# Gene regulatory programs in the life history of *Salpingoeca rosetta*

**DOI:** 10.1101/2023.06.12.544615

**Authors:** Maria Rita Fumagalli, Stefano Zapperi, Caterina AM La Porta

## Abstract

The choanoflagellate *Salpingoeca rosetta* can differentiate into at least five morphologically and behaviorally distinct cell types during its lifetime, going from individual motile cells to linear and rosette-shaped colonies. Due to the capability to form colonies and its close relationship with Metazoa, this organism is considered a model for studying multicellular evolution. The gene regulatory programs underlying these transformations are, however, unknown. Here we analyze transcriptomic data obtained from *Salpingoeca rosetta* in different states to identify a core of genes associated with the formation of multicellular colonies. We then compare the results with other organisms which display simple forms of multicellularity, highlighting commonalities and differences.

## Introduction

The origin of metazoan multicellularity is still a fundamental open intriguing issue in evolutionary biology [1, 2, 3, 4]. The formation of stable colonies of cells is a first fundamental step towards multicellular organisms evolution. A second important and more complex step is the specialization of each cell of the colony in order to divide labor. Hence, it is expected that metazoan multicellularity is linked to basic mechanisms such as cell adhesion, signaling, and differentiation, which could be in common with unicellular and colonial progenitors. Understanding the fundamental gene regulatory processes that may induce transitions from unicellular to multicellular phenotypes is a subject of intense investigation.

Choanoflagellates are aquatic single-cell organisms whose cell morphology displays a single flagellum surrounded by a “collar” of microvilli [5]. The majority of choanoflagellates species are marine although several freshwater species have been described. Due to the close similarity with other metazoan cells, such as the collar cells (choanocytes) of sponges, choanoflagellates are considered to be the closest unicellular relatives of Metazoa [6, 5]. Molecular phylogenetic studies have been used in the past to clarify choanoflagellate evolution and the exact relationship between choanoflagellates and Metazoa [7, 8], firmly established that they are sister groups, sharing a common ancestor.

The ability of choanoflagellates to form colonies has suggested that they could be used as a model system to study the origin of metazoan multicellularity. In particular, the choanoflagellate *Salpingoeca rosetta* (*S. rosetta*) is extremely interesting in this respect, since during its lifetime the same organism can form a linear chains of cells (chain colonies), be found in slow and fast swimmer unicellular states (swimmers), transform into thecate cells (thecate), which are attached to substrates through a secreted structure called theca and finally form a spherical colonies known as rosette [9]. It is thus extremely interesting to study the gene regulatory program underlying these changes and possibly find analogies with similar programs in Metazoa. A detailed study of the transcriptomes of *S. rosetta* showed the involvement of septins in colony formation and indicated that the initiation of colony development may preferentially draw upon genes shared with Metazoa [10]. A later paper showed how the defect in a gene coding for C-type lectin was able to affect the development of the rosette state, showing that this gene was involved in forming an extracellular layer that coats and connects the basal poles of each cell in rosettes [11]. Although adhesion molecules and communication between cells appear to be involved in different type of *S. rosetta* cellular stages [12], it is still unknown if there is a core of genes or a core of pathways which are important for multicellularity.

Here, we tackle this issue by studying differentially expressed genes during the life history of *S. rosetta*, identifying gene-specific signatures and pathways associated with its phenotypes and finally compare with other related organisms. We base our analysis on the gene expression data reported by [10] but focus also on commonalities and differences between distinct colonial forms (i.e rosette and chain), while [10] concentrated on differences between colonial forms and genes representative of single cell. Furthermore, in our analysis we neglect differences due to the presence of single-species or mixed bacterial prey.

## Results and discussion

### Gene expression during *S. rosetta* life hystory

Building on work of [10], we compared gene expression in *S. rosetta* considering two forms of solitary cell types (swimmers and thecate cells, Sw and Th, respectively) and two colonial forms (rosette colony and chain colony, RC and CC, respectively), as described in Materials and Methods section. The four different forms of *S. rosetta* were obtained in the presence of different strains of bacteria, but those have a marginal impact on the gene expression profile [10], as it is shown by the principal component analysis (PCA) reported in Fig.1. Our results suggest that bacteria trigger the transition of single cells toward a colony state mainly by mechanical interactions with *S. rosetta* cells and have a limited effect on gene expression, although we should note that gene expression could also be regulated by post-transcriptional or post-translational mechanisms. Based on our observation, we combined gene expression data obtained in presence of different bacteria. Furthermore, we deviated from the original analysis by [10] by focusing on the specific differences and commonalities between colonial forms (rosette and chain), while [10] concentrated on differences between colonial forms and single cells.

**Figure 1.**
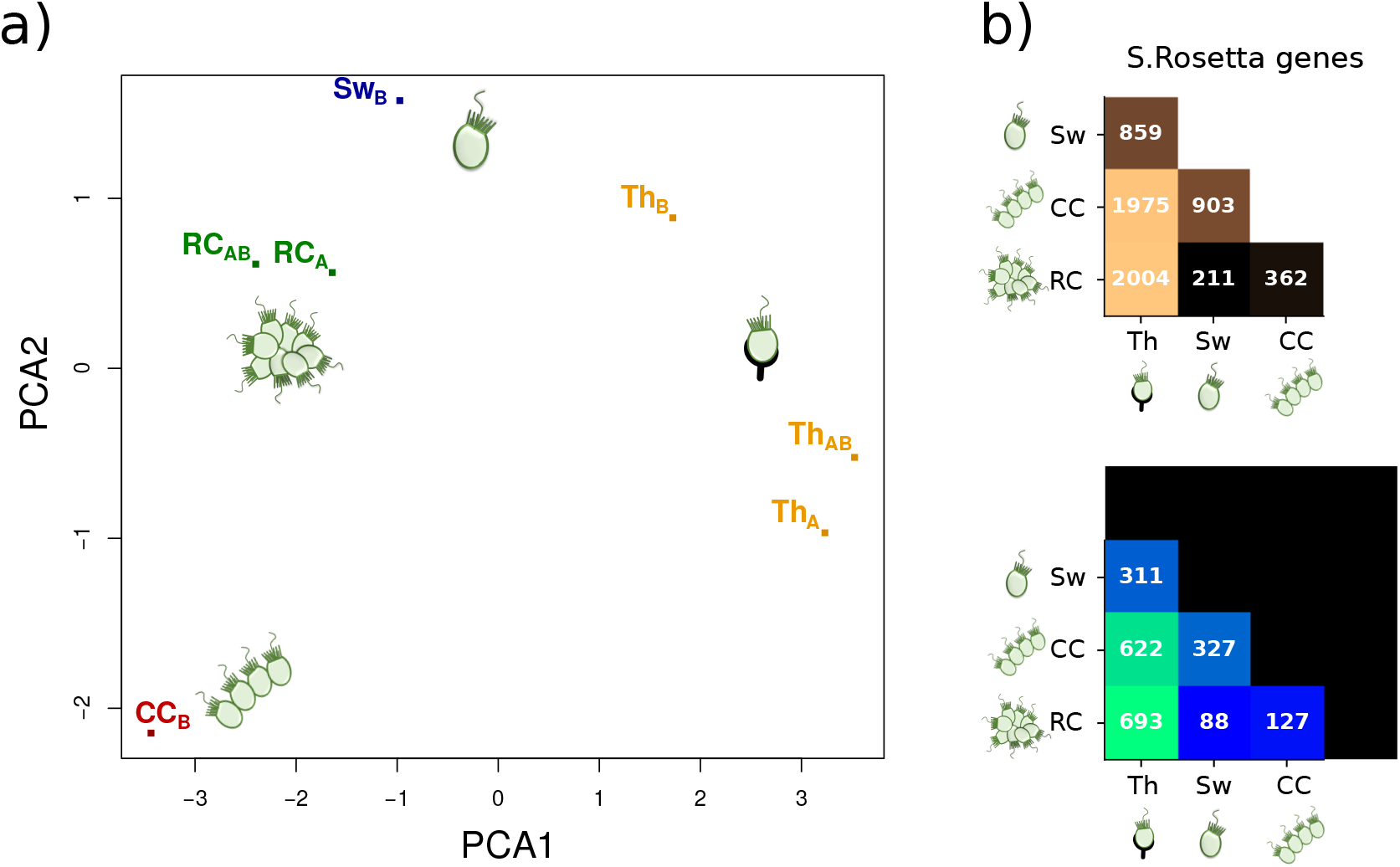
(a) Principal component analysis of gene expression was performed in order to compare all the considered datasets. Data result clustered based on cell type composition, and the presence of bacteria in the growth medium (A. machipongonensis bactera or mixed bacterial prey, as reported by Fairclough et al. (2013). The explained variance for the first and second components is 39% and 21%, respectively. (b) The figure shows the total number of differentially expressed genes between the four cell types for all the *S. rosetta* genes an for the subset mapped on human homologs. Th:thecate cells, Sw:swimmer, CC:Chain Colony, RC:Rosette Colony.

We thus considered gene expression differences across the four cell state of *S. rosetta* (Sw, CC, RC or Th). The first principal component of the PCA allowed us to distinguish between cells forming colonies (CC or RC) and single-cell states (Th and Sw) (see Fig.1a). While Sw cells lay in an intermediate position, there is a strong separation between Th and CC cell states (see Fig.1a). Furthermore, differential expression analysis performed on both the entire set of *S. rosetta* genes and on the subset of putative human orthologues (see Fig.1b top and a bottom, respectively) shows that the Th cell state has the largest number of differentialy expressed (DE) genes when compared to the others states. This seems to be coherent with the fact that the transition from attached cells to chain or colony states requires an intermediate phenotype that might be identified with the swimmer state [9].

Cluster analysis of differentially expressed genes identifies a large group of genes that are downregulated on average from thecate (Th) to chain forming cells (CC) (see red and green clusters in Fig.2). Smaller clusters of genes are slightly downregulated in Th cells when compared to the other three groups (Fig.2b). Taking into account fold changes of single genes, we observe that one third of the genes that are differentially expressed between at least two conditions show a monotonically decreasing expression from Th to Sw cells, and from Sw to colonial cells (CC and RC). Among these, 195 are also significantly dowregulated (FDR <0.05) in CC cells compared to RC state. Interestingly, about a third of these genes are similarly expressed (|log_2_ *FC*| < 1) in Th, Sw and RC cells, suggesting that they are specifically downregulated during the organization of the cells into a chain (CC). On the other hand, only 210 genes show a gradual monotonic increase of expression in the same direction, being highly expressed in CC and poorly expressed in Th cells (Fig.2a), but none of them is significantly differentially expressed between CC and RC cells.

**Figure 2.**
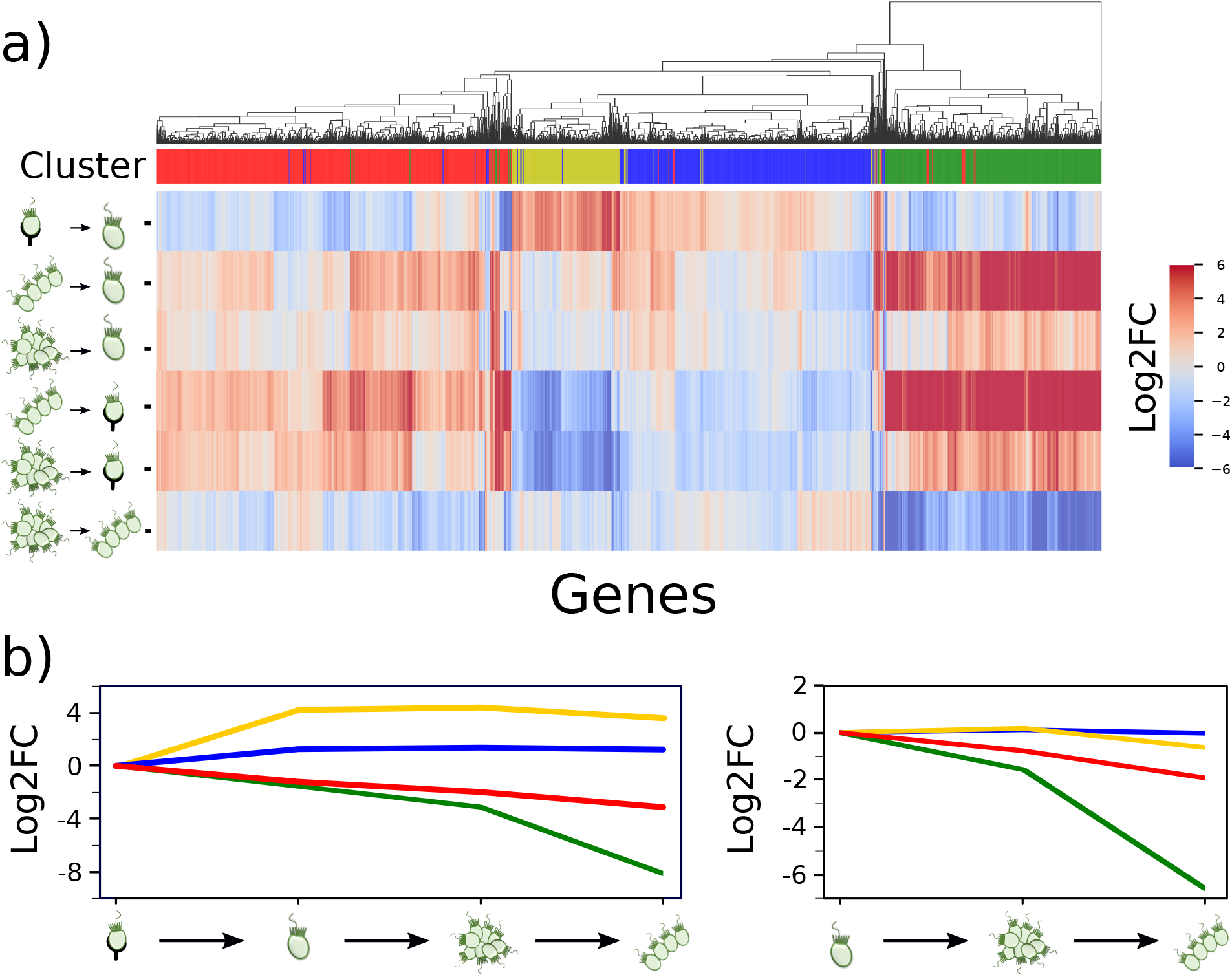
Change of gene expression between solitary cell types (swimmers and thecate cells, SW and Th, respectively) and two colonial forms (rosette and chain, RC and CC, respectively) of *S. rosetta*. a) Heatmap shows the log2FC expression of 2826 genes that are differentially expressed between at least two *S. rosetta* cell types. Hierarchical clustering (dendrogram) and Kmeans clustering (k=4, first row) detected groups of genes having similar expression. b) Plot of the average change of expression of genes belonging to the four clusters using as baseline Th state (left) or Sw (right). Colours as in panel a. Log2FC is calculated between Kmeans clusters centers. A large fraction of genes (clusters red and green) show a coherent regulation from Th to CC cells. Blue cluster include genes slightly downregualted in Th cells compared to all other groups. Yellow cluster contains genes that are upregulated in swimmers and rosette-forming cells compared to Th and CC cells. Heatmaps where obtained in python using the seaborn.clustermap package (v 0.12).

Gene ontology analysis shows that monotonically downregulated genes refer to GO categories such as cell adhesion, cell surface receptor and signaling, ion transmembrane transport and extracellular matrix organization. These genes include cadherins, collagen, protein tyrosine kinase activity receptors and Plexins (Supplementary Table 1). The latter are proteins involved in adhesion processes, synapse development and axon guidance ([13]). Genes that are monotonically upregulated are mainly annotated to GO categories that refer to RNA transcriptional regulation (Supplementary Table 1). A small set of genes participate to response to oxygen-containing compounds including Apolipoprotein-D, LAMTOR5 and alcohol dehydrogenase (Supplementary Table 1). Thus, thecate state does require a strong expression of genes related to extracellular communication with the environment. Cells in colonial state express genes specific of metabolic processes and cell growth regulation.

### Gene specific signature of solitary or colonial *S. rosetta* cell states

We further investigated genes that can be characteristic of the colonial (RC and CC) or solitary cell (Sw or Th) states of *S. rosetta*. Focusing on the differentially expressed genes between these two conditions, we obtained a core of 340 genes with distinct expression in the two conditions (RC and CC) that are expressed at a similar level in both solitary cells (SW and Th, |log *FC*| < 1) but whose expression is at least four fold increased or decreased (|log_2_ *FC*| > 2) in either colony cell state (RC or CC). Among these genes, 106 are in common between the two colony states with a core of 65 genes that are expressed at similar level in either RC or CC cells and 41 genes that are differentially expressed when comparing RC with CC (Fig. 3a, Supplementary Tables 2, 3, 4). These 41 genes include ion-channels, cadherins and serine/threonine kinase activity proteins. There are only 5 genes significantly upregulated in colony cells compared to single cell states. These genes include PTSG 03654 which is annotated as a tyrosine-protein kinase that have no orthologous in other species in OMA but matches to human gene LRRTM2 (p*-*val 10^−24^). The latter is a protein involved in the development and maintenance of excitatory synapse in the vertebrate nervous system [14]. The remaining four genes are PTSG 07401, PTSG 06265, PTSG 07453, PTSG 12548 and two of them are annotated as hydrolase/hydroxylase proteins (Supplementary Table 4). The same analysis has been performed on genes expressed at similar level in single cells (RC and CC) (|log *FC*| < 1) and strongly deregulated (|*log*_2_*FC*| > 2)in comparison to compared to solitary cells (Sw and Th). Fig. 3b shows a small core of 9 genes shared between both solitary cells (Th and Sw) and 6 out of these 9 genes are present also in the core in Fig. 3a. The number of upregulated and downregulated genes for the cases reported in Fig. 3a and Fig. 3b are reported in Fig. 3c and Fig. 3d, respectively. A detailed illustration of the workflow used to identify all the gene groups discussed above, also including the number of upregulated and downregulated genes for each comparison, can be found in Fig. 4.

**Figure 3.**
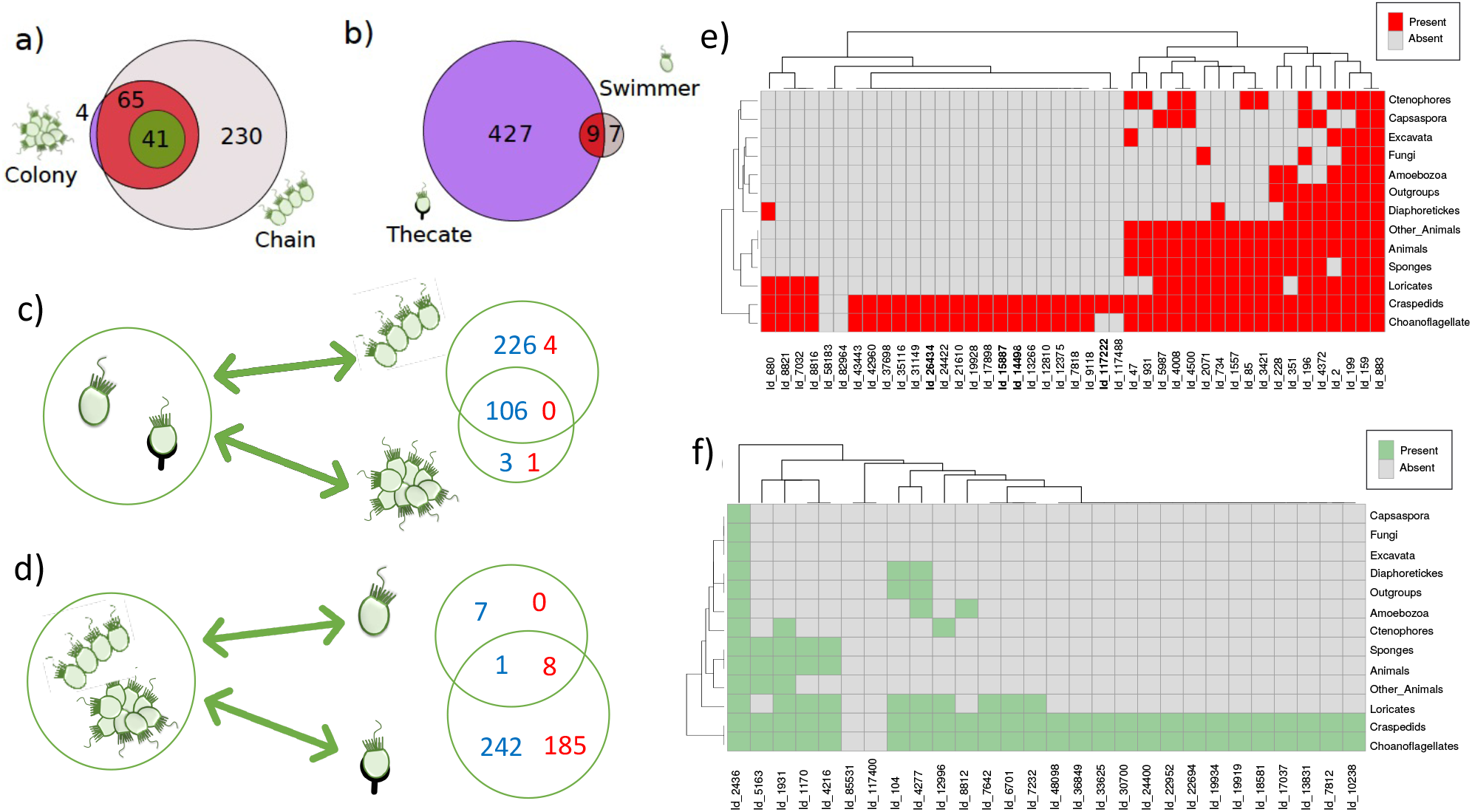
Genes characteristic of single cell and colony states. a) The diagram shows the number of genes differentially expressed in the colony cells (RC and CC) of *S. rosetta* compared to single-cells. We include only those genes not significantly deregulated between Th and Sw (|log *FC*| < 1, *FDR* > 0.05) but having large changes in expression and significant deregulation (|log *FC*| > 2, FDR< 0.05) between these states and either colony or chain. Among these, we selected a core of 65 genes that are differentially expressed between solitary cell states and not differentially expressed between colony cells (RC and CC), while 41 genes are differentially expressed between the two colony states. b) The diagram shows the number of genes not significantly deregulated between colony cells (RC and CC) (|log *FC*| < 1, *FDR* > 0.05) but having large changes in expression and significant deregulation compared to solitary cells (Th and Sw) (|log *FC*| > 2, FDR< 0.05). All the genes shared between solitary states (Th and Sw) are not differentially expressed between the two. c-d) The diagrams display upregulated (red) and downregulated (blue) genes for the cases reported in panels a and b. Up/down regulation refers to the right-side group when compared to the left-side group. e-f) Heatmaps show the degree of conservation of genes specific of single cell or colony state that are expressed at similar level c or differentially expressed d in colony cells (CC and RC). Genes are grouped in families as in and their presence in different classes of organisms is assessed as described in [16]. Names in bold refer to four protein families that include the 6 genes obtained from the intersection of the cores reported in panel a and b (see also main text). Heatmaps where obtained in python using the seaborn.clustermap package (v 0.12).

**Figure 4.**
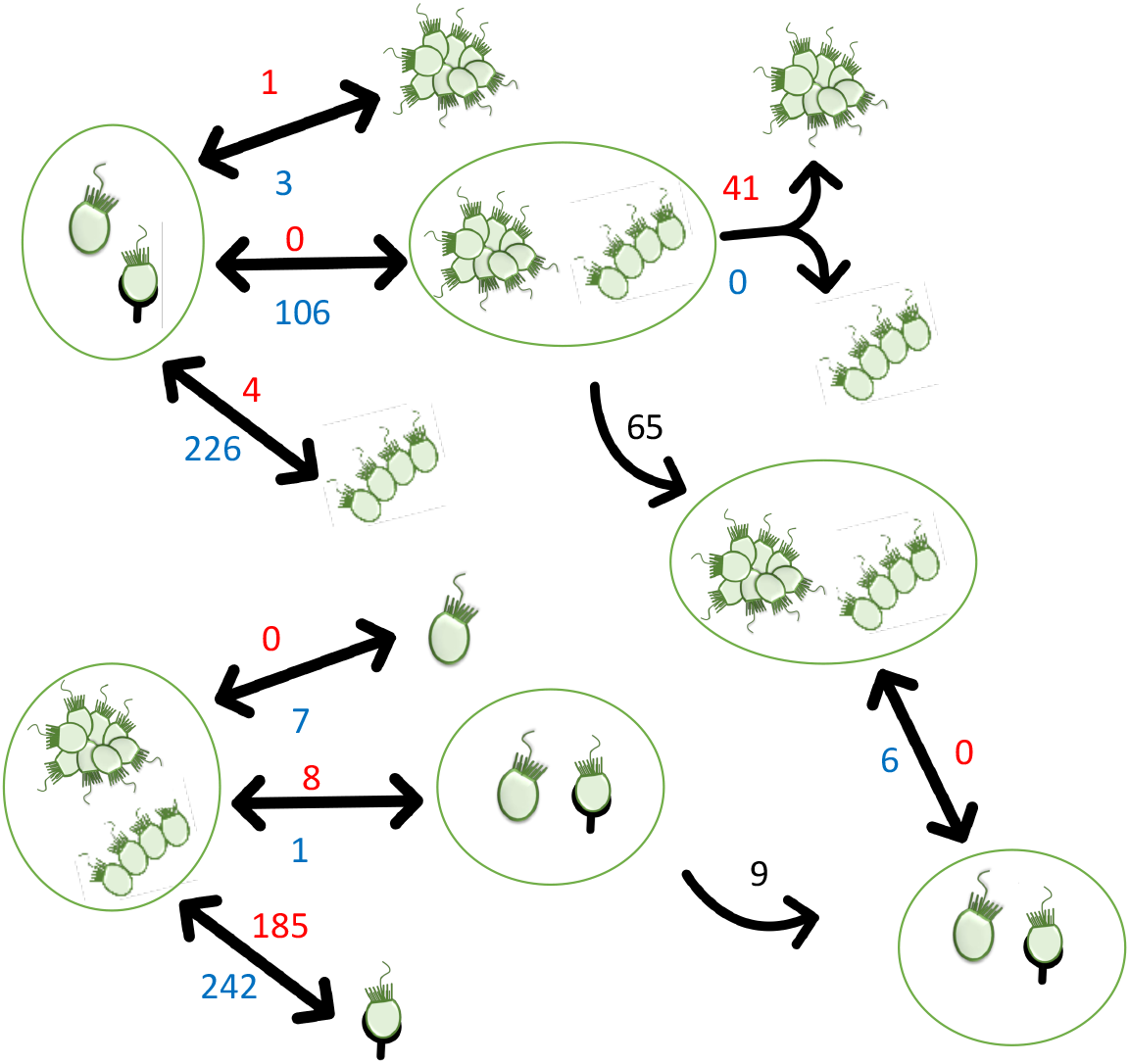
Schematic of the workflow used to identify different gene groups. Double arrows represent differential expression analysis between the groups. Genes that are up/downregulated in the right-side group with respect to the left-side group are shown in red/blue respectively. Single arrows with black numbers represent the number of genes that are expressed at similar level in the two forms enclosed in green circles.

Our results show that there is a very stringent core of genes which includes putative collagen α-1(XII) (PTSG 12412), Javelin-like, isoform A (PTSG 04669), microtubule (MT)-associated protein [15] and protein containing TPR-region (PTSG 12590) that distinguish between solitary and colony cell state (Supplementary Table 5). Four of these genes can be mapped to gene families according to [16] and result to be highly specific of *S. rosetta* and of few other species, being even absent in some choanoflagellates (Fig.3e and Fig. 5). On the other hand, genes that are expressed at similar level in colonial cells (RC and CC) appear to be more conserved among distant species than those distinguishing chain from rosette colonies. These genes are representative of 18 families conserved also in animals (Fig.3e-f and 5) and referred to cadherins, ion channels and other membrane proteins (Supplementary Table 5).

**Figure 5.**
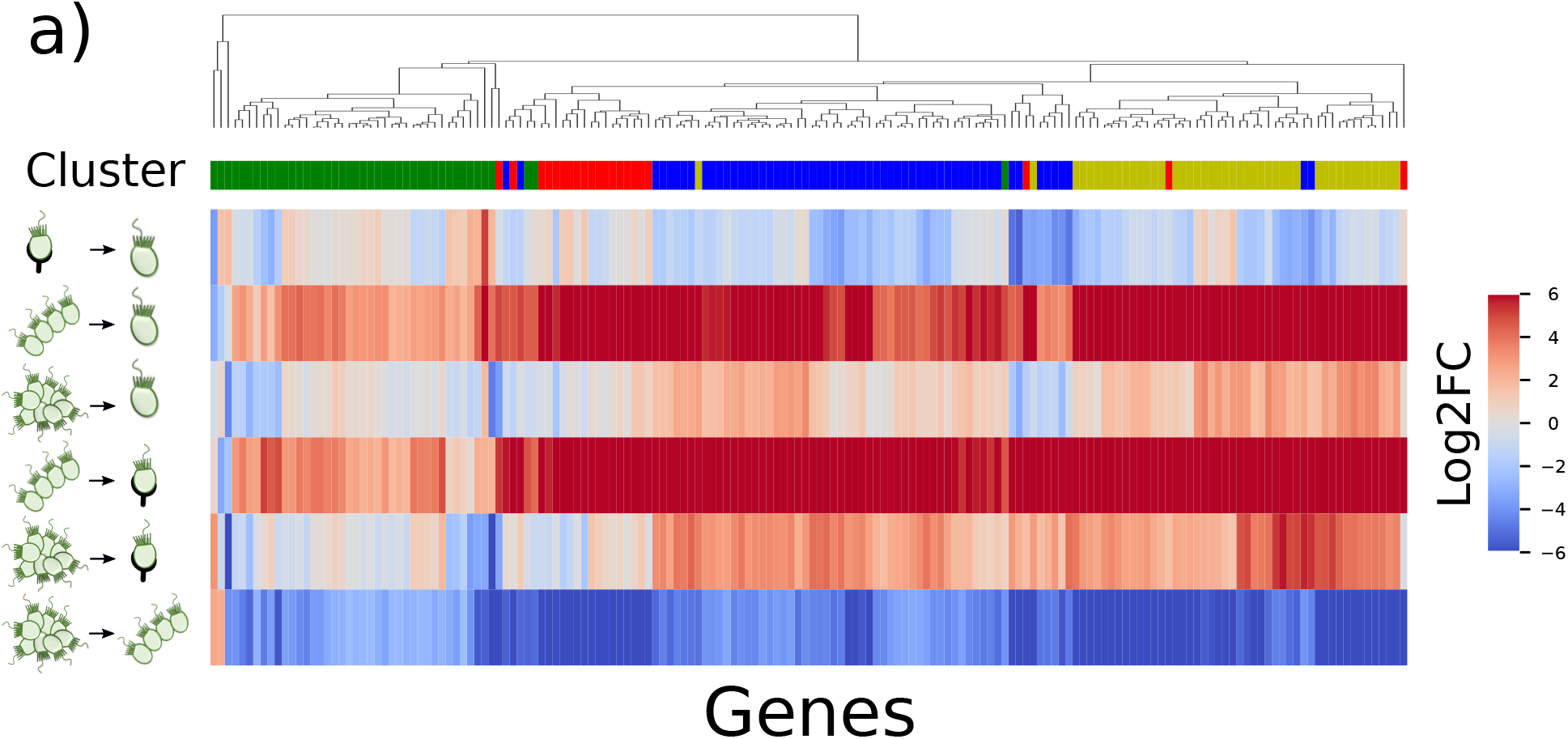
Heatmap shows the log2FC expression of *S. rosetta* human annotated genes that are DE between single and colony cells. Hierarchical clustering (dendrogram) and Kmeans clustering (k=4, first row) detected groups of genes with similar expression. Heatmaps where obtained in python using the seaborn.clustermap package (v 0.12).

### Signature of colonial cell state of *S. rosetta* and comparison with other model organisms

To investigate if the cells in the chain colony are in an intermediate state between solitary cells and cells in the rosette colony or, alternatively, if the chain colony is the result of a completely independent process leading to a polarized layer of specialized cells, we investigated differentially expressed genes between cells belonging to different forms of colony, chain or rosette (with *p* < 0.05 as discussed in the Materials and Methods section). These genes could underlie directional and spatial specific signals, considering that spatial interactions were indeed shown to play a relevant role in the evolution of multicellularity evolution [17]. The analysis of the differentially expressed genes between cells belonging to rosette or chain colonies yields results that are in general very similar to those reported in Fig.2, with a large number of genes that are highly deregulated in chain colony cells (see Fig.6). Overall, about a quarter of the differentially expressed genes are upregulated in chain cells when compared to the other conditions and only 7 genes out of 362 are upregulated in thecate cells when compared to colonial ones. Two of these genes are also conserved in animals, including humans, and annotated as phosphoglycerate dehydrogenase and insulin-induced gene 2 protein.

**Figure 6.**
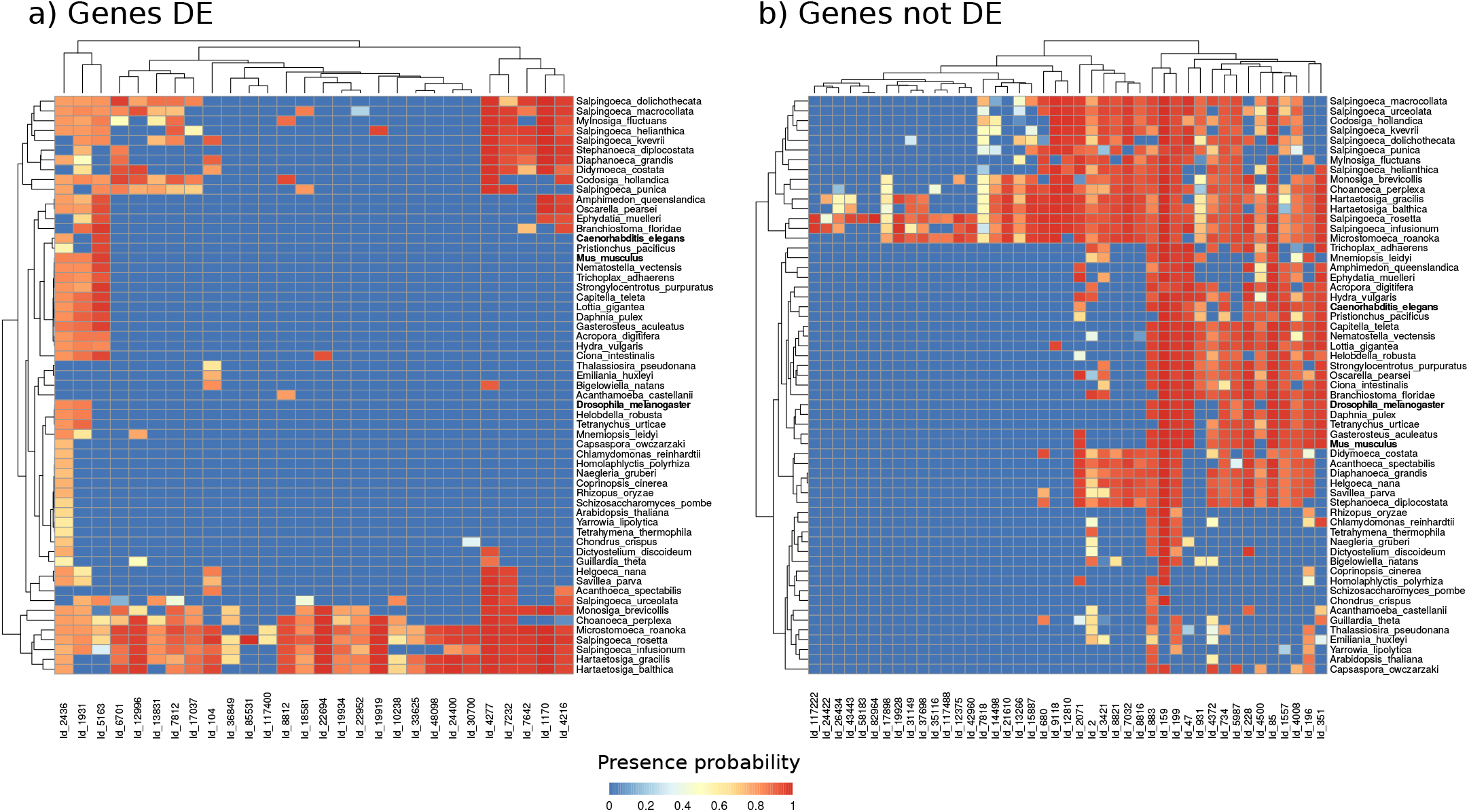
Heatmaps show the degree of conservation of genes specific of single cell or aggregate state that are differentially expressed (a) or expressed at similar level (b) in colony cells (CCand RC). Genes are grouped in families and the probability of presence in different organisms is assessed as described by Richter et al. (2018). Heatmaps where obtained in python using the seaborn.clustermap package (v 0.12).

Figure 3a shows that 41 genes are differentially expressed between RC and CC states even within the core of single-cell specific genes (see also Fig. 4). These genes can be considered to be specific of the rosette colony since they are all downregulated in chain colonies. Functional analysis of the genes (Supplementary Table 6 and 7) results in a wide variety of biological process categories, such as regulation of neurogenesis, cell adhesion and cell migration, that are enriched due to the downregulation in chain colony cells of Plexin-B, Plexin-A4, Plexin-A1, macrophage-stimulating protein receptor, Zinc Finger protein 3 and Smaug Protein, which in human acts as a translational repressor of SRE-containing messengers [18]. Moreover, as shown in figure 3f, we found 5 gene families that are conserved in animals and that contain 7 genes differentially expressed between rosette and chain cells. These include, besides plexins and macrophage-stimulating protein receptor, transgelin (a protein related to microtubules) and RAS guanine-nucleotide exchange factor.

The Rosetteless genes (PTSG 03555, XP 004995286), necessary for colony formation [11], are sightly downregulated in chain colony cells when compared to rosette colony cells (*log*_2_*FC* ≈ −0.34). The genes Jumble (PTSG 00436) and Couscous (PTSG 07368), which are relevant for colony formation of *S. rosetta* cells [19], are slightly upregulated in chain colony cells but Jumble is almost constant in swimmer and rosette colony cells (*log*_2_*FC* ≈ −0.02). The fact that rosetteles genes are critical for rosette formation does not imply a change in their expression. Furthermore, total protein expression or localization are not considered in the RNA sequencing analysis. In any case, data on RNA level of expression can only indirectly related to function: Thus even if the RNA expression levels display small changes, it is conceivable that different localization of the final protein could induce a different shape in the colony.

We finally analyzed the expression pattern of genes related to cell adhesion and related proteins and we compared *S. rosetta* with others organisms such as *V. carteri, C. owczarzaki* and *C. fragrantissima. V. carteri* is a colonial alga that is used as model organism for research into the evolution of multicellularity and organismal complexity [20]. *C. owczarzaki* is an unicellular eukaryotic cell that, similarly to *S. rosetta*, is a close unicellular relative of animals. *C. owczarzaki* assumes three different cellular states during its lifetime. Cells can swim as single cells, form a multicellular aggregates or attach to a substrate [21]. Finally, *C. fragrantissima* is a single-celled protist proposed as a model organism for metazoan multicellularity [22]. Hence, we compared organisms showing multiple states, in order to understand the possible role of adhesion molecules and related functions in the organization of the multicellular state.

We investigated the presence of genes that are putative orthologues of *S. rosetta* genes that are characteristic of multicellular states in other organisms. We find that that the number of pairwise orthologues of *S. rosetta* genes are 2701 for *C. owczarzaki*, 2396 for *C. fragrantissima* and 2224 for *V. carteri*. As shown in Fig. 7a, 983 *S. rosetta* genes have putative orthologues in all the other three species. We next search for the orthologues among the 106 members of the core of genes characteristic of colony state in *S. rosetta*. We find that among these core genes 7 have orthologues in *C. owczarzaki*, 7 in *V. carteri*, and 1 in *C. fragrantissima*. None of these genes have orthologues in all the other three species. The gene having an orthologue in *C. fragrantissima* also has an orhtologue in *V. carteri*. Finally, out of the 7 genes that have orthologues in *C. owczarzaki* only one has an orthologue also in *V. carteri*.

**Figure 7.**
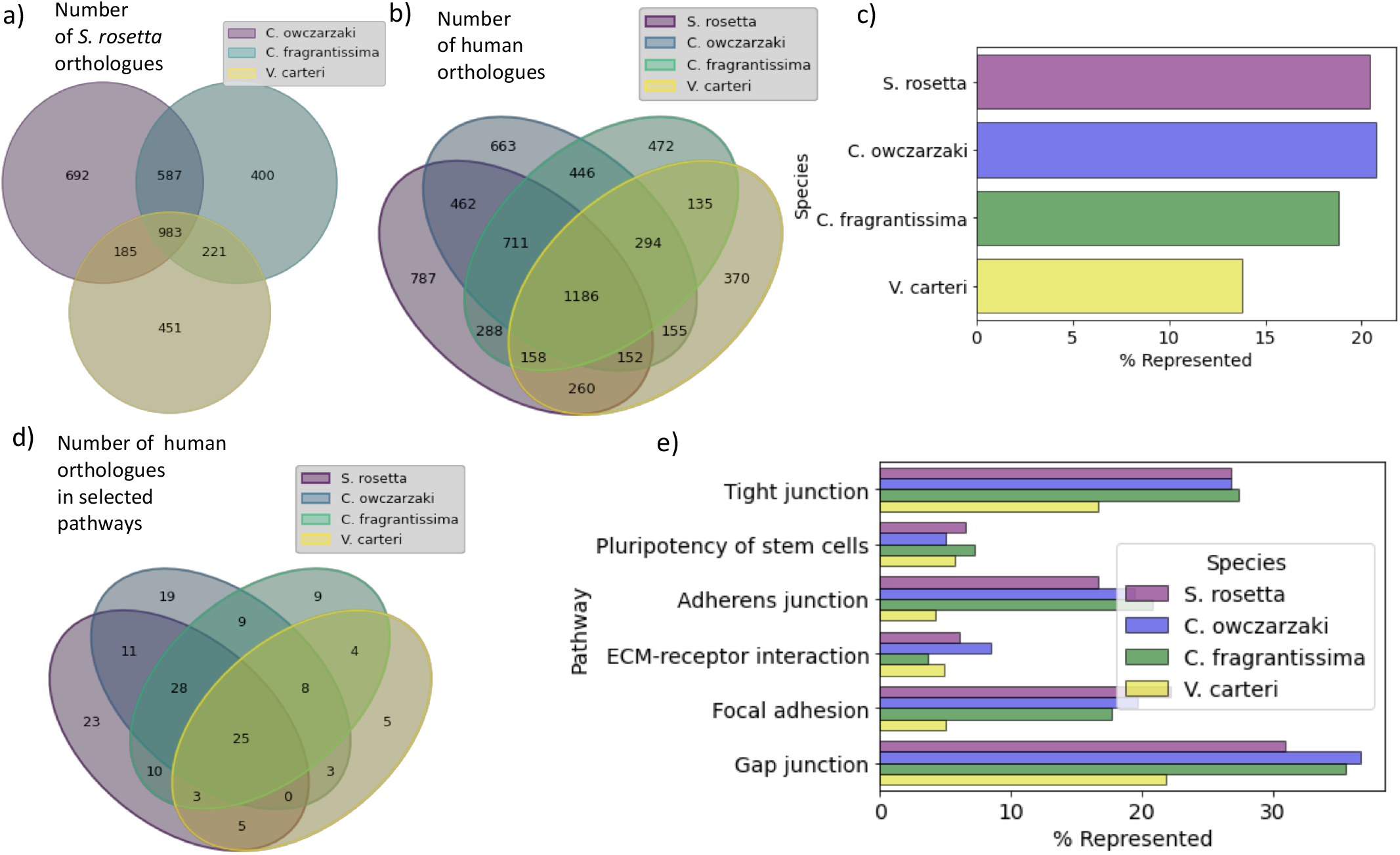
Comparison between *S. rosetta, C. owczarzaki, C. fragrantissima* and *V. carteri*. a) The diagram shows the number of putative orthologues of *S. rosetta* genes in *C. owczarzaki, C. fragrantissima*, and *V. carteri*. b) The diagram shows the number of putative orthologues of human genes in *S. rosetta, C. owczarzaki C. fragrantissima*, and *V. carteri*. c) The percentage of human genes that have an orthologue in the four species considered. d) Same as b but restricted to human genes included in a set of Kegg pathways related to cell-cell and cell-substrate adhesion (i.e. hsa04512 ECM-receptor interaction; hsa04550 Signaling pathways regulating pluripotency of stem cells; hsa04520 Adherens junction; hsa04510 Focal adhesion; hsa04530 Tight junction; hsa04540 Gap junction). e) The percentage of human genes in the pathways that have a putative orthologue in each of the four species considered.

In the case of *C. owczarzaki*, the 7 orthologues of *S. rosetta* core genes are differentially expressed between multicellular and single-cell state in *C. owczarzaki*, even if the expression in the multicellular state is often higher than expression in floating or attached cells. Among these genes, only three are annotated in Uniprot, two as zinc finger proteins and one as ATPase potassium transporter (Supplementary Table 8). However, it is worth recalling that in *C. owczarzaki* attached cells are dramatically different from techate cells in *S. rosetta*, being in a filopodial stage and they transit directly to colony state, without intermediate swimmer state [23]. Thus, we do not expect a net direction of increase/decrease of genes expression from one stage to another. To compare the distance between the genome of the four species and the human genome, we first identify the genes of these species that have a putative human orthologue, as described in the Materials and Methods section. The diagram reported in Fig. 7b shows that there is a group of 1186 genes that have putative orthologues with all the species considered. In general, the number of human orthologues differs among the four species. This is summarized in Fig. 7c reporting the percentage of human gene that are represented in each of the four species. We then considered the genes that can be mapped on human KEGG pathways belonging to cellular community processes and ECM-receptor interaction pathways. The diagram reported in Fig. 7d summarizes the intersection of these genes, showing a core of 25 human genes belonging to the union of the considered pathways that have an orthologue in all the four species. Fig. 7e shows the percentage of genes in each pathway that have an orthologue in each of the pathways. A direct comparison between Fig. 7c and Fig. 7e shows that some pathways such as ECM-receptor interactions are underrepresented while others are overepresented in a way that is largely independent on the species considered.

We also compared the expression level of the *S. rosetta* genes mapped on the selected human KEGG pathways. Out of 340 genes that are differentially expressed between thecate and swimmer with respect to colony or chain, only 20 have a human orthologue and only one of these human orthologues (i.e. Guanylate cyclase soluble subunit beta-1) is included in the selected pathways. Therefore, less than 6% of the differentially expressed genes have a human orthologue and when we consider that in general 39% of *S. rosetta* genes have a human orthologue, we conclude that the gene regulatory program leading to multicellularity in *S. rosetta* is quite distinct from the regulation of adhesion in humans.

## Conclusion

In this paper, we have analyzed the changes in gene expression during the life evolution of *S. rosetta* as the organism transforms from the unicellular swimming state into colony forms. In this way, we identified a core set of genes that are differentially expressed when the cells are either in chain or rosette colony states. It was also possible to identify a second subset of genes that have been used to distinguish cells in rosette from those in chain. This may suggest a specific role of these genes in determining the spatial organization of the colony states. These genes do not appear to be conserved in distant multicellular species, while are present in other unicellular organisms capable to form colonies, such as the members of the *Hartaetosiga* family [8]. We then associated this set of genes to relevant pathways and biological processes and then compare them with genes in related organisms. In particular, we focused on genes related to cell adhesion and highlight commonalities between the formation of colonies in *S. rosetta* and in in another unicellular species, closely related to Metazoa and capable to form transient multicellular colonies. The results we presented suggest that the mechanisms regulating the transition from unicellularity to multicellularity in *S. rosetta* are distinct from those controlling cellular adhesion in humans. We observed, however, gene expression differences between the unicellular state and the thecate state in *S. rosetta*. It would be interesting to investigate further if these changes could be interpreted as evolutionary anticipatory differences related to the cell-extracellular matrix interactions obserbed in Metazoa [24].

## Materials and Methods

### Datasets

*Salpingoeca rosetta, Volvox carteri, Creolimax fragrantissima* and *Capsaspora owczarzaki* fasta protein sequences and corresponding annotation files were downloaded from NCBI website[25]. OMA standalone [26] was used to infer pairwise orthologous between *H. sapiens, S. rosetta, C. owczarzaki, V. carteri, C. reinhardtii, C. fragrantissima* and *S. cerevisiae* (strain ATCC 204508 / S288c). Gene families, group presences, and probabilities of presence in individual species were obtained from Richter et al. [16] for 21 chanoflagellates and 21 other species including animals.

*S. rosetta* gene expression data were retrieved from [10] from *S. rosetta* culture enriched for specific cell-types. Briefly, they sequenced the transcriptome of *S. rosetta* colonial cells forming rosette grown in the presence of A. machipongonensis alone (RCA) or with mixed bacterial prey (RCAB), solitary swimming or chain cells grown in the presence of mixed bacterial prey (SwB, CCB) and single attached cell (thecate) grown with either A. machipongonensis, mixed bacterial prey or a combination of the two (ThA,ThB,ThAB) ([10]). Note that, in this data, it is not possible to distinguish between fast and slow swimmers. List of *C. owczarzaki* genes differentially expressed between adherent, floating and aggregated cells during lifestage was obtained from Sebé-Pedrós and coworkers [23] and downloaded from Figshare repository.

### Differential expression analysis

Differential expression analysis was performed using EdgeR ([27], R package version 3.12.1), implementing default Benjamini-Hochberg correction and genes with corrected p-values < 0.05 were considered as deregulated. For each cellular state (colony, swimmer, chain or single attached) we analyzed separately the different medium condition as well as data averaged over the different growth conditions. We compared the different samples using euclidean distances between gene expression profiles that approximate the typical log2 fold changes between the samples according to Limma package [28].

### Pathway analysis

The list of human genes involved in ECM-receptor interaction, signaling pathways regulating pluripotency of stem cells, adherens junction, focal adhesion, tight junction and gap junction pathways (hsa04512, hsa04550, hsa04520 hsa04510,hsa04530, hsa04540) were downloaded from Kegg Pathways database [29]. None of these pathways involving eukaryotes cellular community processes are represented in Kegg [29] for *S. rosetta, V. carteri* and *C. owczarzaki*. One node of the pathway is considered as active if at least one of the genes mapped on the node is expressed. Panther[30] was used for Pathway and Gene Ontology annotation of *S. rosetta* and human genes.

### Data visualization

Heatmaps in Figs. 2, 3 5 and 6 where obtained in python using the seaborn.clustermap package (v 0.12) available at (https://seaborn.pydata.org/generated/seaborn.clustermap.html)

## Data availability statement

All the data analyzed for this manuscript are publicly available and can be accessed as detailed in the Materials and Methods section.

## Supplementary information

### Supplementary table captions

Supplementary Table 1. Gene ontology and pathway enrichment analysis of genes monotonically up and downregulated from thecate cells to swimmers, rosette and chain-forming cells.

Supplementary Table 2. Gene ontology and pathway enrichment analysis of genes characteristic of RC and CC state and not differentially expressed between rosette and chain-forming cells.

Supplementary Table 3 Complete pathway analysis of genes characteristic of RC and CC state, as reported in Figure 3.

Supplementary Table 4. Complete annotation of genes characteristic of either RC or CC state, as reported in Figure 3.

Supplementary Table 5. Complete annotation of genes characteristic of colonial states and conserved in Animals.

Supplementary Table 6 Annotation and functional enrichment of genes differentially expressed between RC and CC state and characteristic of colonial states.

Supplementary Table 7 Annotation and functional enrichment of all the genes differentially expressed between RC and CC state.

Supplementary Table 8 Differentially expressed genes between single-cell and aggregate states in *S. rosetta* and *C. owczarzaki*

